# Designing of thermostable proteins with a desired melting temperature

**DOI:** 10.1101/2024.09.21.614294

**Authors:** Purva Tijare, Nishant Kumar, Gajendra P. S. Raghava

**Affiliations:** Department of Computational Biology, Indraprastha Institute of Information Technology, Okhla Phase 3, New Delhi-110020, India

**Author notes:** Corresponding Author Prof. Gajendra P. S. Raghava Head and Professor Department of Computational Biology Indraprastha Institute of Information Technology, Delhi Okhla Industrial Estate, Phase III (Near Govind Puri Metro Station) New Delhi, India – 110020 Office: A-302 (R&D Block) Website: http://webs.iiitd.edu.in/raghava/. Equal Contribution (Purva Tijare, Nishant Kumar). Mailing Address of Authors Purva Tijare (PT), Nishant Kumar (NK), Gajendra P. S. Raghava (GPSR).

**Keywords:** Melting Temperature, Prediction, machine learning, Embeddings, Protein Language Models, Thermostable proteins

## Abstract

1.

The stability of proteins at higher temperatures is crucial for its functionality that is measured by their melting temperature (Tm). The Tm is the temperature at which 50% of the protein loses its native structure and activity. Existing methods for predicting Tm have two major limitations: first, they are often trained on redundant proteins, and second, they do not allow users to design proteins with the desired Tm. To address these limitations, we developed a regression method for predicting the Tm value of proteins using 17,312 non-redundant proteins, where no two proteins are more than 40% similar. We used 80% of the data for training and testing; remaining 20% of the data for validation. Initially, we developed a machine learning model using standard features from protein sequences. Our best model, developed using Shannon entropy for all residues, achieved the highest Pearson correlation of 0.80 with an R² of 0.63 between the predicted and actual Tm of proteins on the validation dataset. Next, we fine-tuned large language models (e.g., ProtBert, ProtGPT2, ProtT5) on our training dataset and generated embeddings. These embeddings have been used for developing machine learning models. Our best model, developed using ProtBert embeddings, achieved a maximum correlation of 0.89 with an R² of 0.80 on the validation dataset. Finally, we developed an ensemble method that combines standard protein features and embeddings. One of the aims of the study is to assist the scientific community in the design of targeted melting temperatures. We created a user-friendly web server and a python package for predicting and designing thermostable proteins. Our standalone software can be used to screen thermostable proteins in genomes and metagenomes. We demonstrated the application of PPTstab in identifying thermostable proteins in different organisms from their genomes, the model and data is available at: https://webs.iiitd.edu.in/raghava/pptstab.

**Highlights:**

- Prediction of melting temperature (Tm) on non-redundant proteins
- Machine learning models based on sequence composition and ProtBert embeddings
- A Webserver for predicting Tm and designing thermostable proteins

## 2. Introduction

Proteins are highly versatile organic molecules essential to living organisms, playing a pivotal role in regulating key bodily functions and their ability to carry out the desired functions heavily depends on their thermal stability. The thermal stability of proteins is commonly characterized by their melting temperature (Tm). The melting temperature is a good indicator of the thermal stability of proteins. In drug discovery, the melting temperature is used as a fundamental property. An understanding of factors governing stability is important for the design of stable proteins [1–3]. Industrial production, and pharmaceutical development has broad applications and usage of thermostable proteins [4]. Additionally, thermostable proteins have applications in various fields such as medical research and therapy [5,6]. The experimental techniques used for the identification of protein Tm involve advanced methods such as Mass Spectrometry-based Thermal Proteome Profiling (TPP), Fourier Transform Infrared Spectroscopy, Circular Dichroism, and Differential Scanning Calorimetry [3,7]. These experimental techniques cannot be used to screen thermostable proteins at genome scale due to cost and complexity. There is a need to develop in silico methods that can be used to screen thermostable proteins in protein databases.

In the past, several computational methods have been developed to predict melting temperature of proteins, most of them trained on the redundant datasets shown in Table 1. These methods may fail on proteins which does not have high similarity with proteins in the training dataset. Thus, there is a need to develop a method on a non-redundant dataset of proteins using standard protocols of bioinformatics. In this study, we obtained the dataset from the DeepSTABp, which contains 35,114 protein sequences [4]. We applied the CD-hit at 40% to remove the redundant sequences and got 17,312 non-redundant proteins, with no two protein sequences having more than 40% similarity among them. The data has been used for training, testing and evaluation of our models developed using various machine learning, large language models, and deep learning algorithms. In order to generate features of proteins, we used standard software Pfeature and large language models. The final model was tested on an independent or validation dataset after all the models were trained and tested using 5-fold cross-validation. We developed a web based platform called PPTstab for users to predict Tm of proteins. Design module of PPTstab generates all possible variants of a protein and their Tm, so that users can select a variant with desired Tm. In addition standalone software have been developed to screen thermostable proteins at genome or meta genome scale.

**Table 1:** List of available methods with dataset description and their performance.

| Method | Year | Methodology |
| --- | --- | --- |
| **ProtStab** [8] | 2019 | Gradient boosting of regression trees algorithm with 100 features from PROFEAT, ProtDCal, and ProtParam. |
| **iStable 2.0** [9] | 2020 | Weka and XGBoost for classification and regression models with 10-fold cross-validation. |
| **SCMTPP** [10] | 2021 | A predictor based on support vector machines for the identification and characterization of thermophilic proteins utilizing estimated propensity scores of dipeptides. |
| **ProtStab2** [11] | 2022 | LightGBM algorithm with 6,395 Features; including prothr, ProtDCal, and ProtParam descriptors. |
| **TMPpred** [12] | 2022 | SVM-based thermophilic protein predictor that uses ANOVA on an 188-dimensional feature set to classify thermophilic and non-thermophilic proteins. |
| **SAPPHIRE** [13] | 2022 | A predictor based on a stacking model has been developed to effectively identify thermophilic proteins. |
| **DeepTP** [14] | 2023 | The CNN combined with a Bi-LSTM model is utilized to extract features with long-range dependencies, which are subsequently weighted through a self-attention mechanism and predicted using a Multi-Layer Perceptron (MLP). |
| **BertThermo** [15] | 2023 | Classification of thermophilic proteins using BERT-bfd features and feature engineering methods like SMOTE, LGBM and LR as a classifier. |
| **DeepSTABp** [16] | 2023 | Transformer based PLM for sequence embeddings generation and used as features with MLP as predictor. |
| **DeepTM** [4] | 2023 | A prediction model utilizing Graph Convolutional Neural Networks (GCN), Self-Attention Networks, and Multi-Layer Perceptrons, designed for sequence lengths of fewer than 1028. |
| **ProLaTherm** [17] | 2023 | PLM-based thermophilicity predictor using ProtT5-XL-UniRef50 encoder using sequence information. |
| **TemStaPro** [18] | 2024 | A binary classifier with embeddings generated from ESM-2 and ProtT5-XL protein language models (PLMs). |

## 3. Material and Methods

### 3.1. Main Dataset Acquisition

We have collected 35,114 protein IDs from the DeepSTABp dataset, which was primarily derived from the Meltome Atlas [19]. Their corresponding protein sequences were extracted from the UniProt and UniParc databases, it comprises a wide range of species, including Escherichia coli, Bacillus subtilis, Oleispira antarctica, Thermus thermophilus, Pyrococcus torridus, Geobacillus stearothermophilus, Mus musculus, Homo sapiens (K562 and hepat), Saccharomyces cerevisiae, Caenorhabditis elegans, Danio rerio, Arabidopsis thaliana, and Drosophila melanogaster [16,19,20]. We removed sequences having more than 2500 amino acids and sequences containing non-natural amino acids (e.g., U, Z, O, B, J, X). We employed CD-HIT [21] at 40%, a widely used greedy incremental algorithm for creating a dataset of non-redundant proteins. We ensured that no two sequences had more than 40% similarity and this process yielded a collection of 17,312 unique protein sequences. After that, to maintain fairness in model training and evaluation, we partitioned the dataset into an 80:20 ratio and obtained 13,849 training sequences and 3,463 validation sequences.

### 3.2. Computation of amino acid composition

Amino acid composition, post-translational modifications, protein-protein interaction, and other molecules such as ligands can influence thermal stability in proteins [6,22]. To check the abundance of each amino acid in our datasets, we calculated the composition of an individual amino acid using equation 1 and to know how common or less-common specific amino acids are. First we have divided each of the datasets in positive and negative, followed by calculating the composition of both sets in all the datasets using amino acid composition module of Pfeature software.

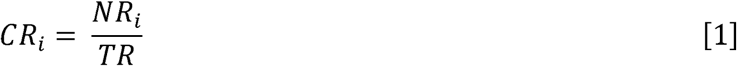

Where, CRi denotes the composition of residue i; NRi indicates the total count of residues of type i; and TR refers to the overall total of residues.

### 3.3. Features Engineering Methods

#### 3.3.1. Composition Based Feature Vectors

For representing the sequence as a numerical vector, we implemented the composition module of Pfeature software [23]. It is a library that calculates features from sequences as well as the structure of the proteins and peptides. It includes five different modules for computing the features: a) Composition-based, b) Binary-profile based, c) Evolutionary information based, d) Structure-based and e) Pattern-based. Utilizing Pfeature, we calculated a diverse array of composition-based features, encompassing 15 distinct types, including amino acid composition (AAC), residue repeat information (RRI), distance distribution of residues (DDOR), atomic composition (ATC), bond composition (BTC), composition based on physico-chemical properties (PCP), conjoint triad calculation (CTC), composition enhanced transition and distribution (CeTD), Shannon entropy of the entire protein (SEP), Shannon entropy for all residues (SER), Shannon entropy derived from physico-chemical properties (SPC), quasi-sequence order (QSO), sequence order coupling number (SOCN), pseudo amino acid composition (PAAC), and amphiphilic pseudo amino acid composition (APAAC). The vector matrix of these descriptors is presented in Table 2.

**Table 2:** Description of the features with their vector matrix extracted using Pfeature.

| Type of Feature | Vector Size |
| --- | --- |
| AAC | 20 |
| APAAC | 23 |
| ATC | 5 |
| BTC | 4 |
| CeTD | 189 |
| CTC | 343 |
| DDR | 20 |
| PAAC | 21 |
| PCP | 30 |
| QSO | 42 |
| RRI | 20 |
| SEP | 1 |
| SER | 20 |
| SOCN | 2 |
| SPC | 25 |

#### 3.3.2. Embeddings Feature Vectors

We created sequence embeddings utilizing ProtBert, a pre-trained protein language model (PLM) that is founded on the BERT natural language processing algorithm [24,25]. This model was trained on the Big Fantastic Database (BFD), which comprises more than 2.3 million protein sequences. From the final hidden layers of ProtBert, we obtained 1024-dimensional vectors that serve as the representations of protein sequences, commonly referred to as embeddings. The main distinction between this model and the original Bert model lies in its approach to sequence handling; unlike the original Bert, which utilizes next-sentence prediction technique, ProtBert treats each sequence as a complete document. It employs a random masking style for up to 15% of amino acids as input. Our methodology involves training through the tokenization of sequences composed of uppercase amino acids, with a single space separating each token. The vocabulary consists of 20 distinct tokens, each representing the linear structures of the 20 standard amino acids (A, C, D, E, F, G, H, I, K, L, M, N, P, Q, R, S, T, W, Y, V), while non-natural amino acids are not included. For optimization, we employed Lamb; a layerwise adaptive large batch optimization technique with a learning rate set at 0.002 and a weight decay parameter of 0.01. This approach yields an embedding vector of 1024 dimensions, situating each sequence within a high-dimensional space to encapsulate the nuanced relationships among amino acids by analyzing the context of their occurrence [26].

Additionally, we extracted embeddings using various LLMs such as Ankh, ProtGPT2, ProtT5- XL-Uniref50, and ProsT5. The Ankh model represents a pioneering general-purpose protein language model, developed utilizing Google’s TPU-v4, featuring a reduced parameter count of less than 10% for pre-training, under 7% for inference, and below 30% for the embedding dimension [27]. ProtGPT2, on the other hand, is a decoder-only transformer model that has undergone pre-training on the UniRef50 protein database (version 2021_04) with a causal modeling objective, employing the GPT2 transformer architecture. This model comprises 36 layers and has a dimensionality of 1280, amounting to a total of 738 million parameters [28]. ProtTrans, particularly ProtT5-XL-U50, is derived from the T5-3b model, which is a text-to-text Transfer Transformer (T5) pre-trained on an extensive collection of protein sequences in a self-supervised manner, utilizing raw protein sequences without any human annotations to automatically generate inputs and labels. Unlike the original T5-3b, which used a span denoising objective, ProtT5-XL-UniRef50 employed a BART-like masked language model (MLM) denoising objective [25]. ProsT5 aka Protein Structure-Sequence T5 is a bilingual protein language model designed to translate between protein sequences and structures, developed by fine-tuning ProtT5-XL-U50 with a dataset of 17 million proteins and high-quality 3D structure predictions from AlphaFoldDB [29].

### 3.4. Model Development

We started by evaluating multiple algorithms to construct a robust predictor. For this process, we utilized H2O AutoML, a scalable automatic machine learning library, version 3.46.0.4 [30]. We applied all available regression algorithms to our training dataset and evaluated their performance using R^2^ and PCC regression-based metrics. From the evaluations, we chose the top 5 performing models, including Support Vector Regression (SVR), Extra Trees Regressor (ET), Nu Support Vector Regression (NuSVR), Light Gradient Boosting Machine (LGBM), and Multi- layer Perceptron (MLP). These algorithms were then implemented in our private notebooks using the scikit-learn library for Python, while LGBM was implemented in the standard LightGBM implementation [31,32]. Throughout our evaluation, we implemented them using default parameters and aimed to identify the model that demonstrates the highest predictive capability for our specific task.

Ultimately, we created a custom Artificial Neural Network (ANN) model using TensorFlow, consisting of four dense hidden layers with 256, 128, 64, and 32 units, activated by a ReLU (rectified linear unit), followed by an output layer [33]. The model is trained using means squared error loss and the Adam optimizer to prevent overfitting [34]. Furthermore, the custom ANN model and the MLP regressor model was integrated to create a unified predictive framework, enabling a comprehensive evaluation of their collective effectiveness in prediction.

### 3.5. Cross Validation

For model evaluation, we have implemented the K-fold cross-validation (CV) technique, a widely used method for assessing model performance and ensuring robustness. This technique involves dividing the entire dataset into k subsets or folds. In our study, we select k=10, resulting in ten distinct folds where each fold represents a portion of the dataset. The k-1 folds are used for training the model, while the remaining folds are used for validation. Each time the process is repeated, the validation set is the same. The model is evaluated on the one validation fold after training on the k-1 training folds. This cyclic process ensures that each fold is tested, providing a comprehensive assessment of the model’s performance across different segments of the data. After completing all iterations, we calculate the mean of the performance metrics from each fold. This average serves as the overall performance measure of the model, offering a robust estimate of its generalization ability and reducing the risk of overfitting [35].

### 3.6. Normalization

Normalization is a crucial technique used in data preprocessing to ensure that all data in a dataset fall within a similar range. This uniformity is essential as it enhances the performance of many machine learning models. Some models may not be optimal without it. The Min-Max Scaler (MMS) method scales down the data to fall within a range of [0, 1]) or [-1, 1]), making it more suitable for model training and analysis [36]. The Min-Max Scaler achieves this by applying a mathematical transformation to the data, ensuring that all features are proportionally adjusted relative to each other. Mathematically, the procedure is calculated using the equation 2.

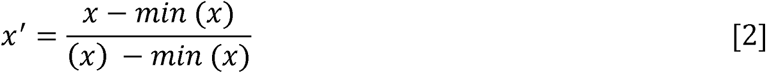

### 3.7. Performance Assessment Metrics

To assess the effectiveness of our regression model, we computed five standard evaluation metrics. These metrics are essential for assessing the predictive capability of regression models across various dimensions.

**Coefficient of determination (R^2^):** The proportion of variance in the dependent variable that is predictable from the independent variable is called the coefficients of determination. R2 estimates how close the data is to the regression line. The closer the value is to 1 the better the regression model is.

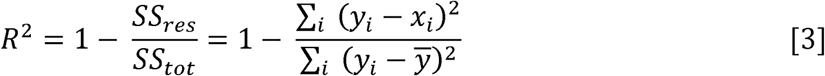

SS_tot_ is the total sum of squares, SS_res_ is the sum of squares of residuals. yi is the true value and xi is the prediction.

**Mean squared error (MSE):** The mean squared error is the average of the squares of the errors. The difference between the estimated and actual values is called the average squared difference.

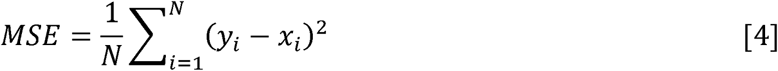

A sample of N data points is used to generate a N prediction, with the *xi* observed values of the *yi* variable being predicted.

**Mean average error (MAE):** MAE indicates the error between paired values for predictions and observations.

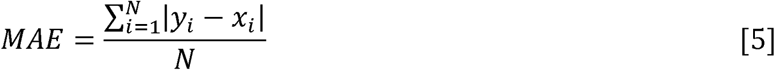

where yi is the prediction and xi is the true value.

**Root mean squared error (RMSE):** The RMSE is the measure of the differences between the predicted and observed values.

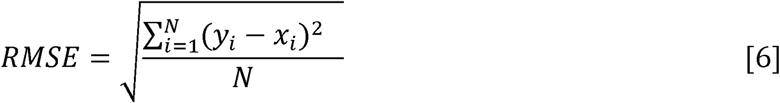

The predicted value is yi, the experimental value is xi.

**Pearson correlation coefficient (PCC):** The Pearson correlation coefficient quantifies the relationship between two data sets. A normalized measurement of covariance is the difference between the product of two variables’ standard deviations. The range is -1 to 1 and is represented in a mathematical way as:

Here, σ_X_ is the standard deviation of X, σ_Y_ is the standard deviation of Y, μ_X_ is the mean of X, μ_Y_ is the mean of Y and E is the expectation [37].

## 4. Results

The result section has five different categories including: (i) Data analysis of Thermostable Proteins, (ii) Machine Learning methods, (iii) Hybrid approaches, (iv) Web Server and (v) Application. The complete workflow of the study is illustrated in Figure 1, and the details of the following subsections can be found below.

**Figure 1:**
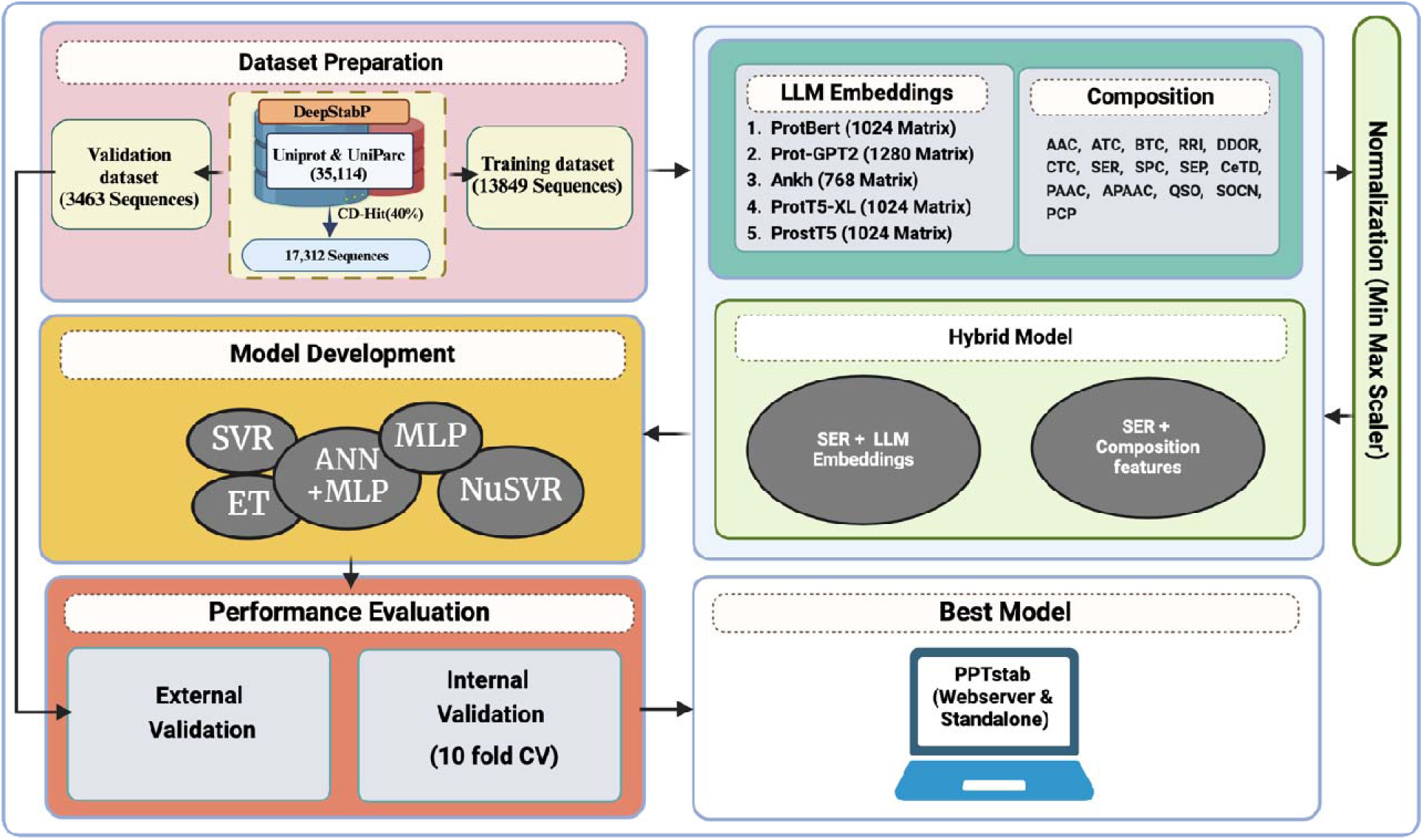
Overview of the study’s complete workflow

### 4.1. Data Analysis

We performed compositional and correlation analysis on the main dataset to understand the relationship between Tm values and the composition of residues.

#### 4.1.1. Composition Analysis of Proteins

Ponnuswamy et al. tested the relationship between the amino acid composition and thermal stability of globular proteins [38]. Our proteins were divided into two groups; proteins with Tm > 50°C and protein with Tm < 50°C, then average amino acid composition was calculated for each residue in both of the groups of proteins as shown in Figure 2. From the figure; we could see that in thermophilic proteins Leucine (L), Alanine (A), Glycine (G), and Glutamic Acid (E) are significantly abundant, whereas Serine(S), Lysine(K), Glutamine(Q) and Histidine(H) are mainly found in proteins with Tm < 50°C.

**Figure 2:**
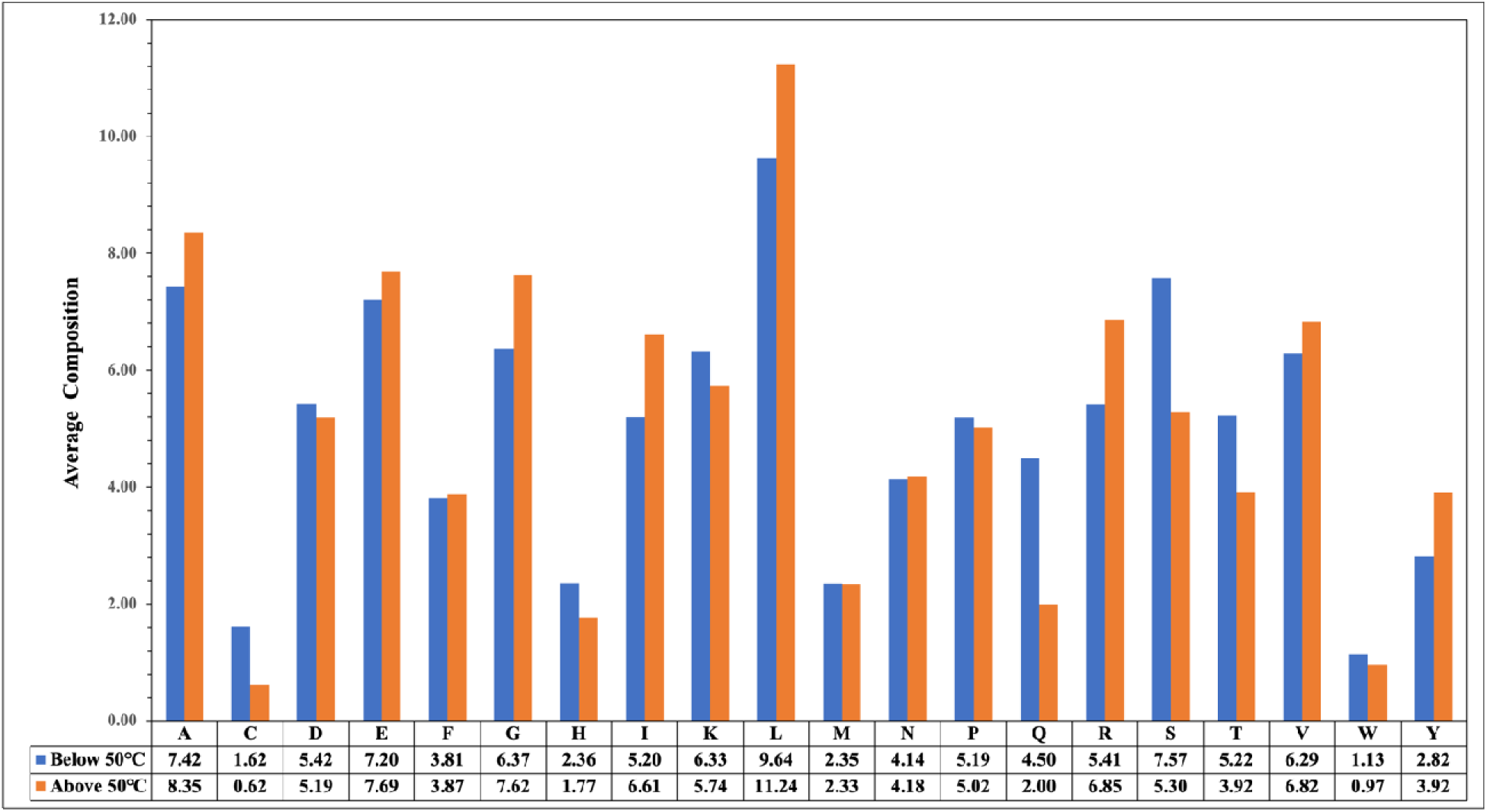
Average percent amino acid composition of thermostable proteins

#### 4.1.2. Correlation between Tm and descriptors

We first computed the percent composition of each descriptor, including AAC, SER, PAAC, and APAAC, that achieved the maximum performance, as shown in Table 4. Secondly, we have computed the correlation between the composition of a residue and Tm for each type of residue shown in Supplementary Figures S1, S2, S3, and S4. The two most highly positive and negative amino acids are shown in Table 3, which lists the top correlations for different compositions.

**Table 3:**
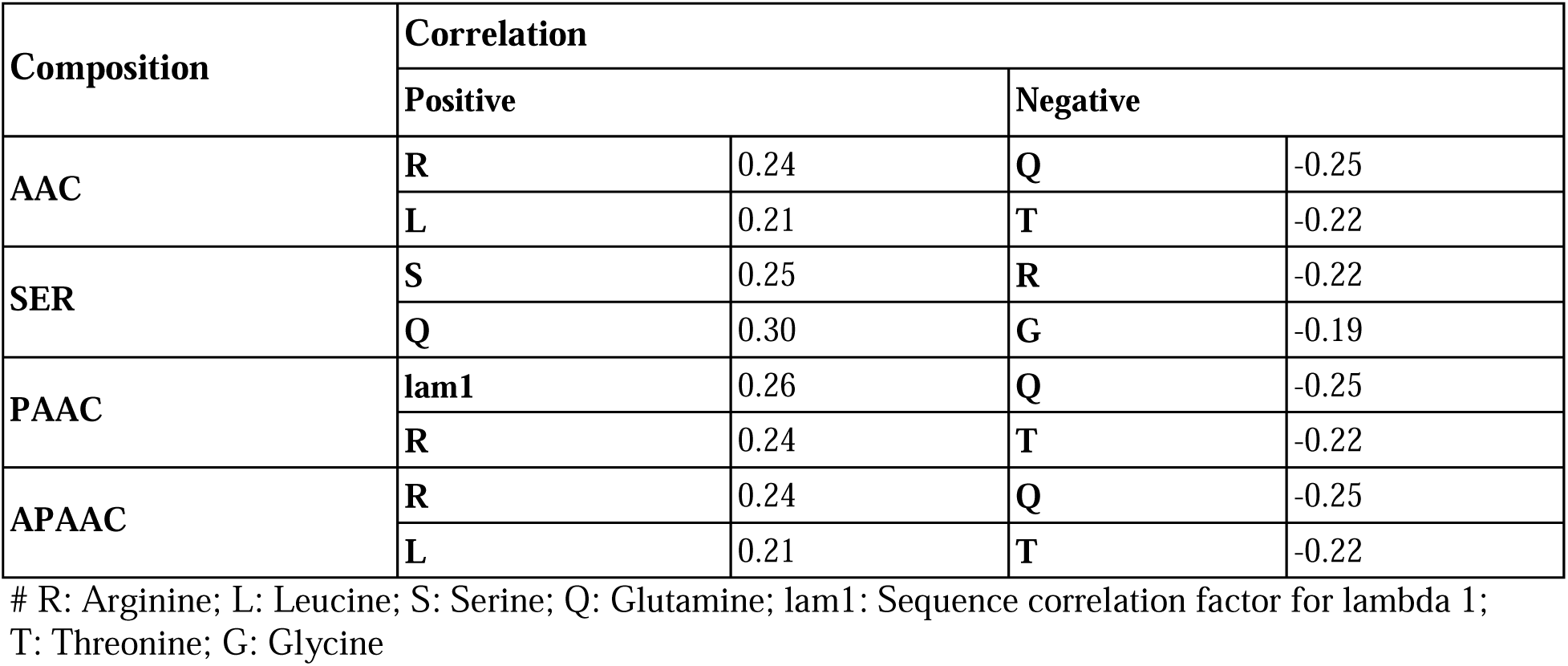
The top positive and negative correlation in different compositions.

| Composition | Correlation |  |  |  |
| --- | --- | --- | --- | --- |
|  | Positive |  | Negative |  |
| AAC | R | 0.24 | Q | -0.25 |
|  | L | 0.21 | T | -0.22 |
| SER | S | 0.25 | R | -0.22 |
|  | Q | 0.30 | G | -0.19 |
| PAAC | lam1 | 0.26 | Q | -0.25 |
|  | R | 0.24 | T | -0.22 |
| APAAC | R | 0.24 | Q | -0.25 |
|  | L | 0.21 | T | -0.22 |

**Table 4:** Performance of ANN+MLP regressor on different composition-based features.

| Name & Description of descriptor | Training |  | Validation |  |
| --- | --- | --- | --- | --- |
|  | PCC | R<sup>2</sup> | PCC | R<sup>2</sup> |
| AAC (Amino acid composition) | 0.81 | 0.65 | 0.77 | 0.51 |
| ATC (Atomic composition) | 0.53 | 0.28 | 0.45 | 0.17 |
| BTC (Bond composition) | 0.29 | 0.08 | 0.26 | 0.07 |
| PCP (Physico-chemical properties based composition) | 0.78 | 0.60 | 0.68 | 0.45 |
| RRI (Residue repeat information) | 0.49 | 0.24 | 0.25 | 0.07 |
| DDR (Distance distribution of residues) | 0.74 | 0.54 | 0.64 | 0.40 |
| SER (Shannon entropy for all residues) | 0.80 | 0.64 | 0.80 | 0.63 |
| SEP (Shannon entropy based on physico-chemical properties) | 0.50 | 0.25 | 0.47 | 0.22 |
| CTC (Conjoint triad calculation) | 0.63 | 0.38 | 0.66 | 0.42 |
| PAAC (Pseudo amino acid composition) | 0.81 | 0.64 | 0.78 | 0.59 |
| APAAC (Amphiphilic pseudo amino acid composition) | 0.81 | 0.65 | 0.79 | 0.59 |
| QSO (Quasi-sequence order) | 0.80 | 0.64 | 0.63 | 0.32 |
| SOCN (Sequence order coupling number) | 0.34 | 0.11 | 0.32 | 0.10 |
| CETD (Composition enhanced transition distribution) | 0.74 | 0.54 | 0.65 | 0.07 |
| SPC (Shannon Entropy at Property Level) | 0.74 | 0.55 | 0.71 | 0.49 |

### 4.2. Machine Learning Methods

In this study, we have used various machine learning algorithms to build models for temperature prediction including ET, MLP, NuSVR, LGBM, SVR, and ensemble models such as ANN+MLP regressor. We have developed these models using compositional features, LLM embeddings, and hybrid features.

#### 4.2.1 Development of Prediction Models

After developing various algorithms using different compositional features, we found that the ensemble model ANN+MLP outperformed all other regression models. Table 4 presents the performance metrics of this ensemble model across different compositions and descriptors. We observed that among 17 different compositions, features like SER, APAAC, PAAC, and AAC emerged as the primary contributors among others. Notably, the SER feature achieved the highest performance with an R² value of 0.63 and a PCC value of 0.80 on the validation dataset.

For a comprehensive overview of the results with different regressors, please refer to Supplementary Table S1.

#### 4.2.2. Composition-based Models

Based on the above observations, we systematically developed a method using the SER composition of proteins as input features, with each feature represented by a 20-dimensional vector corresponding to the 20 different amino acids [39]. Various models were trained on a specific dataset containing SER descriptors known as the SER to predict the Tm value as the output. The comprehensive results obtained through 10-fold cross-validation with normalization as preprocessing are summarized in Table 5. However, the SER features alone did not yield good results, as seen by the ensemble model achieving the R² value of 0.63 and a pearson correlation coefficient (PCC) of 0.80.

**Table 5:**
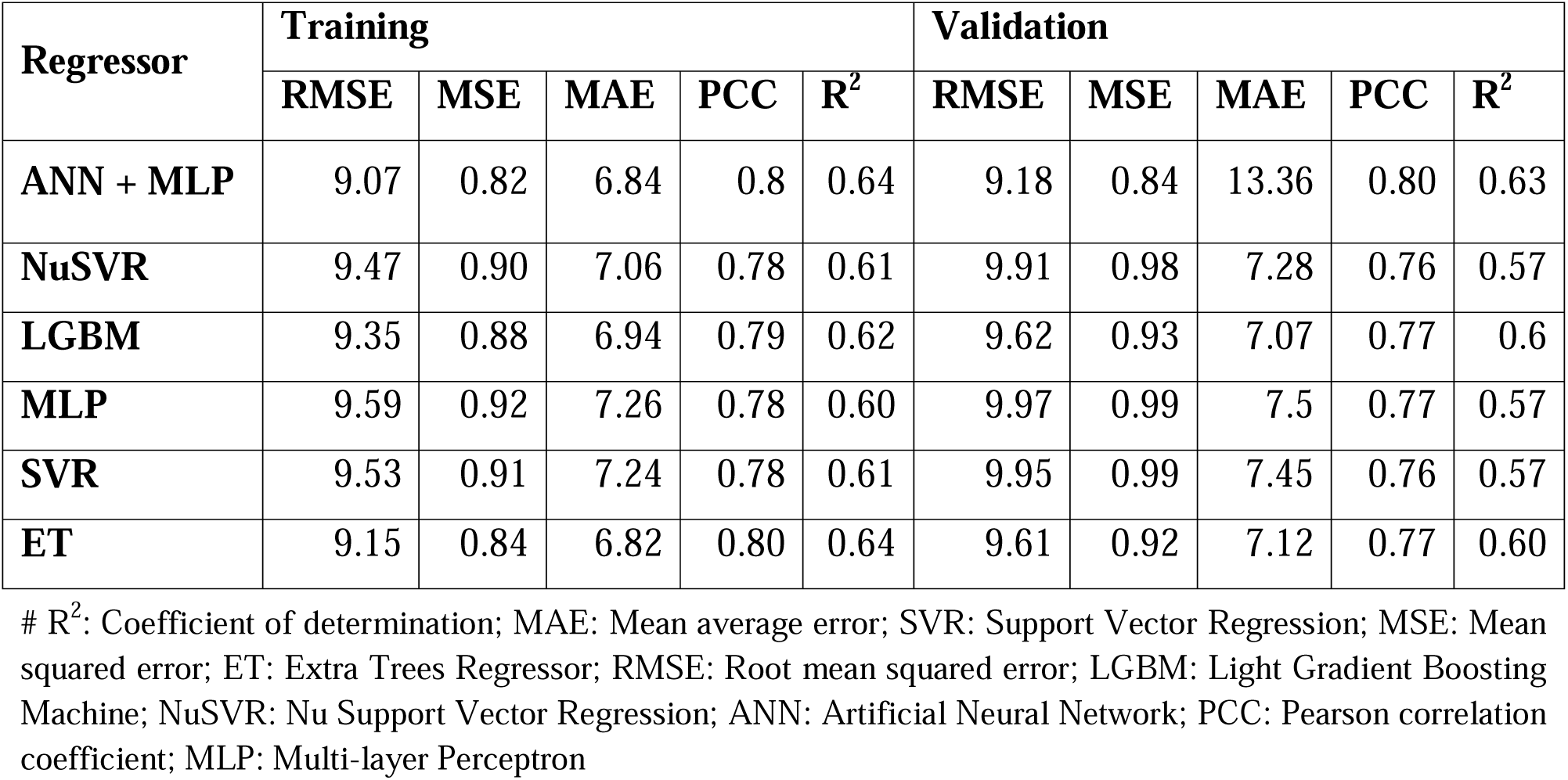
Performance of different regressors on SER features/descriptors.

#### 4.2.3. LLM Embeddings-based Models

In this study, we developed a prediction model using Protein Language Model (PLM) embeddings, specifically using the pre-trained LLM known as ProtBert. It was used as a static feature encoder without any fine-tuning to derive protein embeddings for input, forming the foundation of our method [40]. These embeddings were then combined with categorical flags indicating ’lysate’ or ’cell’ conditions, creating a comprehensive dataset of 1026 features used to train our predictive model for temperature values. The next step involved training various regressors with the aim of enhancing performance. ProtBert embeddings significantly outperformed other embeddings derived from models such as Ankh, ProtGPT2, ProtT5-XL- Uniref50, and ProsT5.

Among the models we evaluated, the ensemble model ANN+MLP proved to be the most effective predictor for Tm values, achieving impressive metrics including a R^2^ value of 0.80 and a PCC value of 0.89. Additionally, the model demonstrated impressive accuracy with a Mean Absolute Error (MAE) of 3.00, Root Mean Squared Error (RMSE) of 4.11, and Mean Squared Error (MSE) of 0.28 as shown in Table 6. These results show a huge decrease in error rates compared to prior methods, highlighting the improved generalizability and robustness of our approach. Additionally, we explored embeddings from different LLMs without fine-tuning and implemented a variety of machine learning algorithms using these embeddings. The detailed performance results of these models are provided in Supplementary Table S2, showcasing the robustness and superiority of the ProtBert embeddings in our predictive framework.

**Table 6:** Performance of different regressors using various LLM embeddings.

| Features | Models | Training |  |  |  |  | Validation |  |  |  |  |
| --- | --- | --- | --- | --- | --- | --- | --- | --- | --- | --- | --- |
|  |  | RMSE | MSE | MAE | PCC | R<sup>2</sup> | RMSE | MSE | MAE | PCC | R<sup>2</sup> |
| ProtBert | ANN+MLP | 3.97 | 0.26 | 3.00 | 0.90 | 0.81 | 4.11 | 0.28 | 3.00 | 0.89 | 0.80 |
|  | CatBoost | 4.46 | 0.33 | 3.32 | 0.87 | 0.76 | 4.58 | 0.35 | 3.34 | 0.86 | 0.75 |
|  | NuSVR | 4.60 | 0.35 | 3.38 | 0.86 | 0.75 | 4.67 | 0.36 | 3.36 | 0.86 | 0.74 |
| ProtT5-XL-UniRef50 | ANN+MLP | 4.07 | 0.28 | 3.07 | 0.89 | 0.80 | 4.12 | 0.28 | 3.09 | 0.89 | 0.79 |
|  | CatBoost | 4.45 | 0.33 | 3.32 | 0.87 | 0.76 | 4.49 | 0.34 | 3.35 | 0.87 | 0.76 |
|  | NuSVR | 4.39 | 0.32 | 3.33 | 0.88 | 0.77 | 4.36 | 0.32 | 3.23 | 0.88 | 0.77 |
| Ankh | ANN+MLP | 4.93 | 0.41 | 3.61 | 0.84 | 0.71 | 5.03 | 0.42 | 3.61 | 0.83 | 0.69 |
|  | CatBoost | 5.88 | 0.58 | 4.35 | 0.77 | 0.59 | 5.90 | 0.58 | 4.30 | 0.76 | 0.58 |
|  | **NuSVR** | 5.88 | 0.58 | 4.38 | 0.77 | 0.59 | 5.89 | 0.58 | 4.36 | 0.76 | 0.58 |
| **ProtGPT2** | **ANN+MLP** | 4.94 | 0.41 | 3.61 | 0.84 | 0.70 | 5.25 | 0.46 | 3.64 | 0.82 | 0.67 |
|  | **CatBoost** | 5.44 | 0.49 | 3.99 | 0.80 | 0.65 | 5.54 | 0.51 | 4.00 | 0.79 | 0.63 |
|  | **NuSVR** | 5.65 | 0.53 | 4.15 | 0.79 | 0.62 | 5.87 | 0.57 | 4.23 | 0.77 | 0.58 |
| **ProstT5** | **ANN+MLP** | 5.33 | 0.48 | 3.88 | 0.81 | 0.66 | 5.24 | 0.46 | 3.77 | 0.82 | 0.67 |
|  | **CatBoost** | 6.58 | 0.72 | 4.83 | 0.69 | 0.48 | 6.62 | 0.73 | 4.79 | 0.69 | 0.47 |
|  | **NuSVR** | 6.10 | 0.62 | 4.54 | 0.74 | 0.55 | 6.12 | 0.63 | 4.50 | 0.74 | 0.55 |

### 4.3. Hybrid Approaches

In this approach, we developed models by combining various feature combinations as input. From our analysis, we observed that embeddings from the ProtBert LLM outperformed other composition-based features. Among the composition-based features, SER showed the best performance with a PCC value of 0.80 and a R² value of 0.63 as shown in Table 4. To further improve the performance, we combined SER descriptors with different feature sets including AAC, PAAC, and ProtBert embeddings. In this approach, we utilized a vector matrix consisting of 1046 features which includes SER and ProtBert embeddings along with the ‘lysate’ or ‘cell’ flag. We have developed models using different regressors, and it was observed that the ensemble model ANN + MLP did not perform well compared to prior approaches, achieving a PCC value of 0.89 and an R^2^ value of 0.79. The complete SER results are presented in Table 7, while the comprehensive results of other features are provided in Supplementary Table S3. After computing the performance of different feature sets, we found that combining SER features with ProtBert embeddings did not achieve greater performance compared to the ProtBert embeddings model.

**Table 7:** Performance of combined feature set: SER descriptors and ProtBert embeddings.

| Models | Training | Validation |
| --- | --- | --- |

|  | **RMSE** | **MSE** | **MAE** | **PCC** | **R<sup>2</sup>** | **RMSE** | **MSE** | **MAE** | **PCC** | **R<sup>2</sup>** |
| --- | --- | --- | --- | --- | --- | --- | --- | --- | --- | --- |
| **ANN + MLP** | 3.99 | 0.27 | 3.02 | 0.90 | 0.80 | 4.05 | 0.27 | 2.97 | 0.89 | 0.79 |
| **NuSVR** | 7.67 | 0.59 | 5.63 | 0.86 | 0.75 | 8.06 | 0.65 | 5.81 | 0.86 | 0.72 |
| **LGBM** | 7.51 | 0.56 | 5.54 | 0.87 | 0.76 | 7.67 | 0.59 | 5.58 | 0.86 | 0.74 |
| **MLP** | 7.23 | 0.52 | 5.49 | 0.88 | 0.77 | 7.51 | 0.56 | 5.62 | 0.88 | 0.76 |
| **SVR** | 7.78 | 0.61 | 5.88 | 0.86 | 0.74 | 8.09 | 0.65 | 6.00 | 0.85 | 0.72 |
| **ET** | 7.46 | 0.56 | 5.50 | 0.87 | 0.76 | 7.59 | 0.58 | 5.51 | 0.87 | 0.75 |

## 5. Web Interface

To assist the scientific community, we developed an easy-to-use and freely available web server called “PPTstab” available at https://webs.iiitd.edu.in/raghava/pptstab/. This tool provides the prediction of thermal melting temperature for protein sequences, using embeddings as features computed by the protein language model named ProtBert. Our best-performing model was integrated within the modules “Predict” and “Design”. The first module employs an ANN+MLP regressor along with composition features derived from protein sequences for prediction, while the second module utilizes embeddings calculated via ProtBert, a powerful LLM available at https://github.com/agemagician/ProtTrans and https://huggingface.co/Rostlab/prot_bert. The web server was developed using a responsive HTML template and is compatible with a number of operating systems. Additionally, to help users easily predict Tm value for their sequence on a larger scale, we built a python-based package named “PPTstab”.

## Application of PPTstab

PPTstab is an advanced tool designed to predict the melting temperature (Tm) and help identify thermostable proteins. Thermostable proteins which are present in a wide array of organisms from extremophiles thriving in harsh conditions to commonly studied model organisms like bacteria, plants, and mammals are useful for various purposes [41,42]. Industrial application of thermostable proteins include enzyme catalysis in biofuel production, food processing, and pharmaceutical manufacturing, where their ability to withstand temperature fluctuations is crucial. Also useful in medical research and therapy, serving as stable components in drug delivery systems, diagnostic assays, and therapeutic agents. Addressing critical clinical needs, such as developing heat-stable vaccines for global distribution and resilient biologics for targeted disease treatments [43].

In our analysis, we utilized reviewed proteomes from the Uniprot database, focusing on organisms adapted to extreme temperature environments. This includes microorganisms, such as psychrophiles, mesophiles and thermophiles, which can thrive at temperatures as low as 0°C, and above 60°C [44]. Among the species analyzed was Psychrobacter frigidicola, a gram negative, non-motile, aerobic and osmotolerant bacteria. Used in biotechnological processes like restriction endonucleases, uracil-DNA glycosylases, and producing bioactive metabolites with medical applications [45]. Shewanella oneidensis is an electrochemically active mesophilic bacterium with maximum growth temperature of ∼35°C, which has been extensively studied for advancing bioelectrochemistry [46]. Lastly we analysed Thermus thermophilus, a well-known thermophile used in biotechnological applications such as genetic manipulation, structural genomics, and systems biology [47,48]. The summary of our findings is showcased in the Table 8.

**Table 8:** Categorical distribution of microorganisms with their melting temperature ranges.

| Category | Micro organism | Count | 30°C-40°C | 40°C-50°C | 50°C-60°C | 60°C-70°C | 70°C-80°C | 80°C-90°C |
| --- | --- | --- | --- | --- | --- | --- | --- | --- |
| Psychrophile | *Psychrobacter frigidicola* | 2298 | 0.26 % | 24.07% | 53.68% | 16.98% | 4.70% | 0.30% |
| Mesophile | *Shewanella oneidensis* | 4068 | 0.32% | 27.24% | 52.19% | 15.71% | 4.40% | 0.15% |
| Thermophile | *Thermus thermophilus* | 2227 | 0.05% | 4.28% | 31.25% | 32.37% | 22.51% | 9.55% |

Here, the percentages indicate how each microorganism is distributed across different temperature ranges. From this table, we learn that Psychrobacter frigidicola shows the most protein stability at 50°C-60°C, indicating its adaptability beyond cold environments. Shewanella oneidensis exhibits broad stability from 40°C-70°C, reflecting its preference for moderate temperatures and Thermus thermophilus has the highest stability at 60°C-70°C, typical of thermophiles. This analysis highlights that while each microorganism is adapted to a specific temperature range, their proteins display stability across a broader range, underscoring evolutionary flexibility. The analysis also showcases the utility of PPTstab in identifying thermostable proteomes, and thereby guiding potential applications in biomedical as well as biotechnological fields [49].

## Discussion and Conclusion

The cost of experimental methods used for identifying the thermostability of proteins are expensive and time-consuming in nature; therefore the computational techniques are a better alternative to predict the melting temperature of proteins. It is crucial for understanding their behavior and functionality across diverse biological processes and applications. In this study, we have introduced a method called “PPTstab” that was developed on a non-redundant dataset, unlike previously available methods that were developed using redundant datasets. Thus, an accurate and efficient prediction method using primary sequences is needed. We obtained the dataset from a previously developed method named ‘DeepSTABp’ that has mostly taken data from the Meltome Atlas study. Then we applied CD-hit (40%) to sequences for reducing redundancy.

Afterward, we applied various machine learning algorithms such as ET, SVR, LGBM, NuSVR, and MLP regressor to different composition-based features and sequence embeddings from the protein language models, along with the ‘lysate’ or ‘cell’ flag as shown in Table 2. We also computed the performance on individual composition descriptors and different LLM model embeddings and merged the composition with LLM embeddings. Among all the composition-based models, SER demonstrated superior performance. Therefore, we have combined the SER features with other features to construct the hybrid model but merging did not achieve good results, so we excluded it. The ensemble model created using ANN with MLP regressor trained on the ProtBert embeddings outperforms every model, achieving an R2 value of 0.80 and a PCC value of 0.89 on the validation dataset, which consists of a 1026-feature vector.

Despite the similar performance of Prot-Bert and ProtT5-XL-UniRef50 models in training and testing scenarios, we chose Prot-Bert due to its lower computational cost and faster results. While LLMs have made significant contributions across various domains, deploying them effectively poses numerous challenges. These include high training and maintenance costs, scalability issues, limited causality understanding, short attention spans, restricted transfer learning capabilities, and gaps in non-textual context understanding. Additionally, LLMs struggle with generating long-form text, collaborating effectively, handling ambiguity, incremental learning, and managing unstructured data and input errors. Fine-tuning LLMs for specific tasks is a useful technique but requires huge amounts of memory and compute resources, thereby limiting its access to only a few institutions. Therefore, computational demand remains a significant barrier for fine-tuning LLMs [50]. We have provided a user-friendly web server PPTstab (https://webs.iiitd.edu.in/raghava/pptstab/) for predicting, and designing thermostable proteins. We hope that this method will significantly contribute to the scientific community working in this domain.

## Authors’ contributions

GPSR collected the dataset. PT, and NK processed the dataset. PT, and GPSR implemented the algorithms and developed the prediction models. NK, PT, and GPSR analysed the results. NK created the front-end and back-end of the webserver. NK, PT, and GPSR penned the manuscript. GPSR conceived and coordinated the project. All authors have read and approved the final manuscript.

## Author’s Biography

1. Purava Tijare is a Project Fellow in Computational Biology at the Department of Computational Biology, Indraprastha Institute of Information Technology, New Delhi, India.
2. Nishant Kumar is currently working as Ph.D. in Computational biology from Department of Computational Biology, Indraprastha Institute of Information Technology, New Delhi, India.
3. Gajendra P. S. Raghava is currently working as Professor and Head of Department of Computational Biology, Indraprastha Institute of Information Technology, New Delhi, India.

## Abbreviations

ANN: Artificial Neural Network
BFD: Big Fantastic Database CV: cross-validation
ET: Extra Trees Regressor
LGBM: Light Gradient Boosting Machine
LLM: Large Language Model
MAE: Mean average error
MLM: masked language model
MLP: Multi-layer Perceptron
MMS: Min-Max Scaler
MSE: Mean squared error
NuSVR: Nu Support Vector Regression
PLM: Protein Language Model
PCC: Pearson correlation coefficient
R^2^: Coefficient of determination
ReLU: rectified linear unit
RMSE: Root mean squared error
SVR: Support Vector Regression
Tm: melting temperature

## Acknowledgments

Authors are thankful to the University Grants Commission (UGC), and Department of Biotechnology (DBT) for fellowships and financial support, and the Department of Computational Biology, IIITD New Delhi for infrastructure and facilities. We would like to acknowledge that Figures were created using BioRender.com.

## Data Availability Statement

All the datasets used in this study are available at the “CLBTope” web server, https://webs.iiitd.edu.in/raghava/pptstab/data.html.

