## Supplementary Figures for "Designing of thermostable proteins with a desired melting temperature"

**Mailing Address of Authors**

**# Equal Contribution**

***Corresponding Author**

Prof. Gajendra P. S. Raghava

Head and Professor

Department of Computational Biology

Indraprastha Institute of Information Technology, Delhi

Okhla Industrial Estate, Phase III (Near Govind Puri Metro Station)

New Delhi, India – 110020 Office: A-302 (R&D Block)

Website: <http://webs.iiitd.edu.in/raghava/>

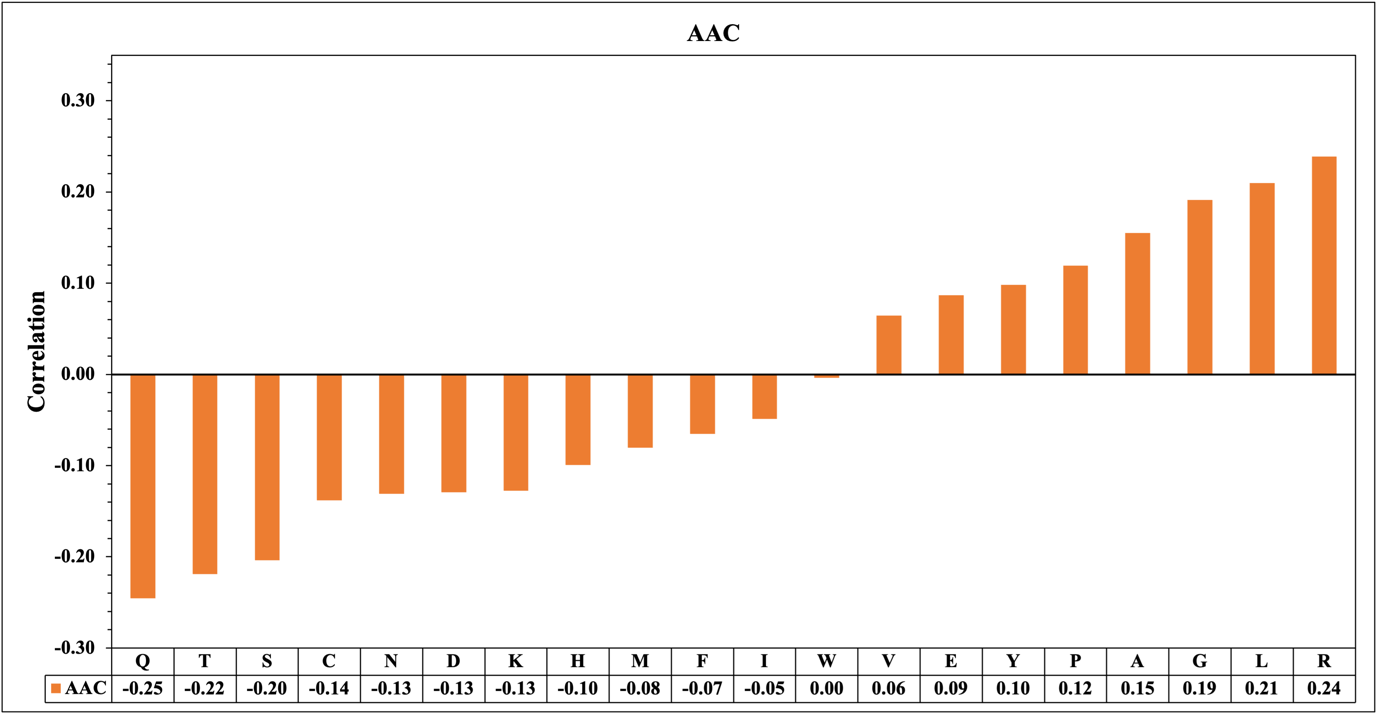

**Supplementary Figure S1: The correlation between the percent composition of AAC residues and Tm**

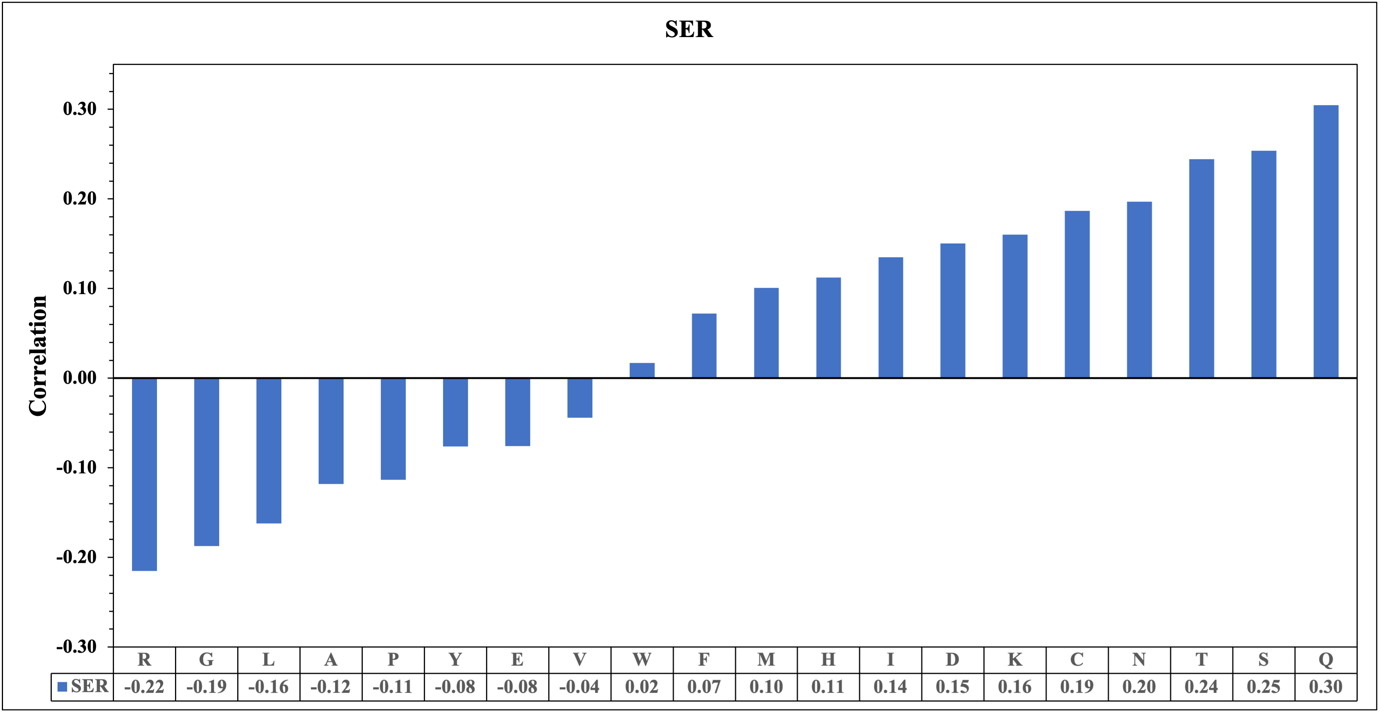

**Supplementary Figure S2: The correlation between the percent composition of SER residues and Tm**

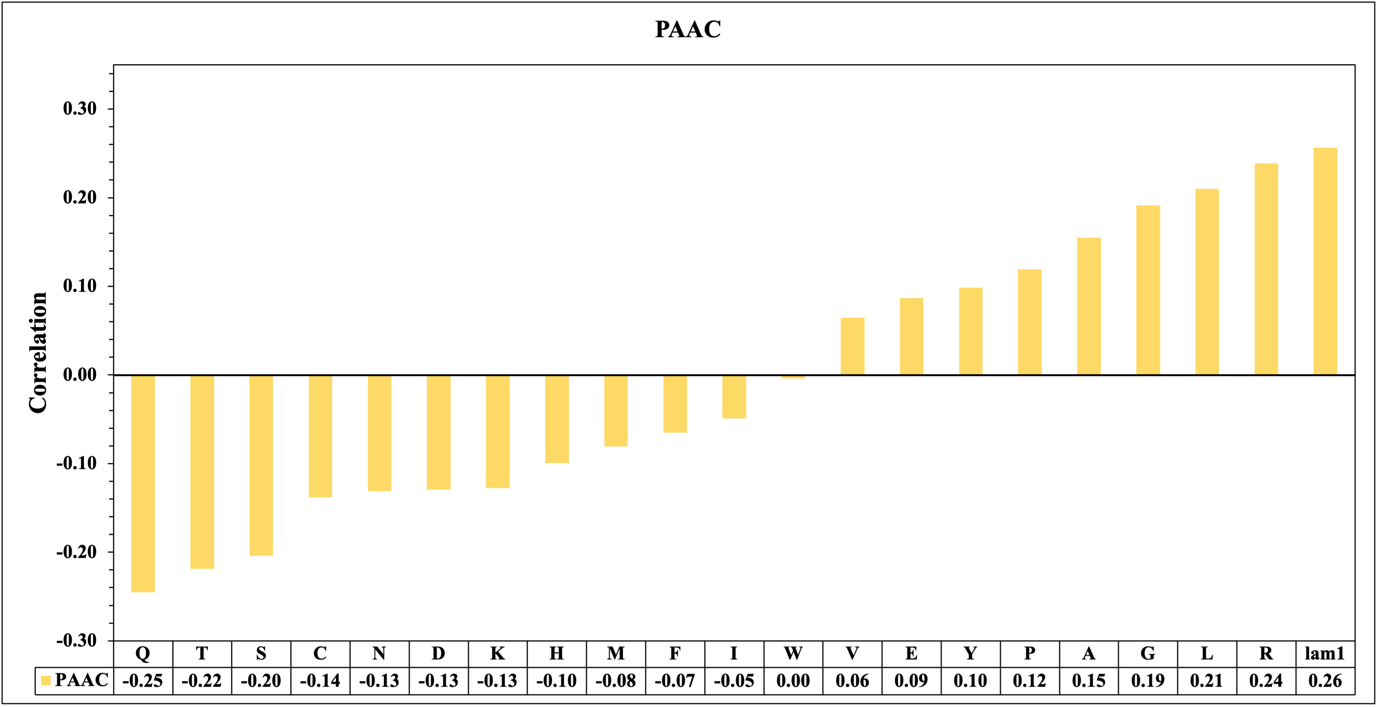

**Supplementary Figure S3: The correlation between the percent composition of PAAC residues and Tm**

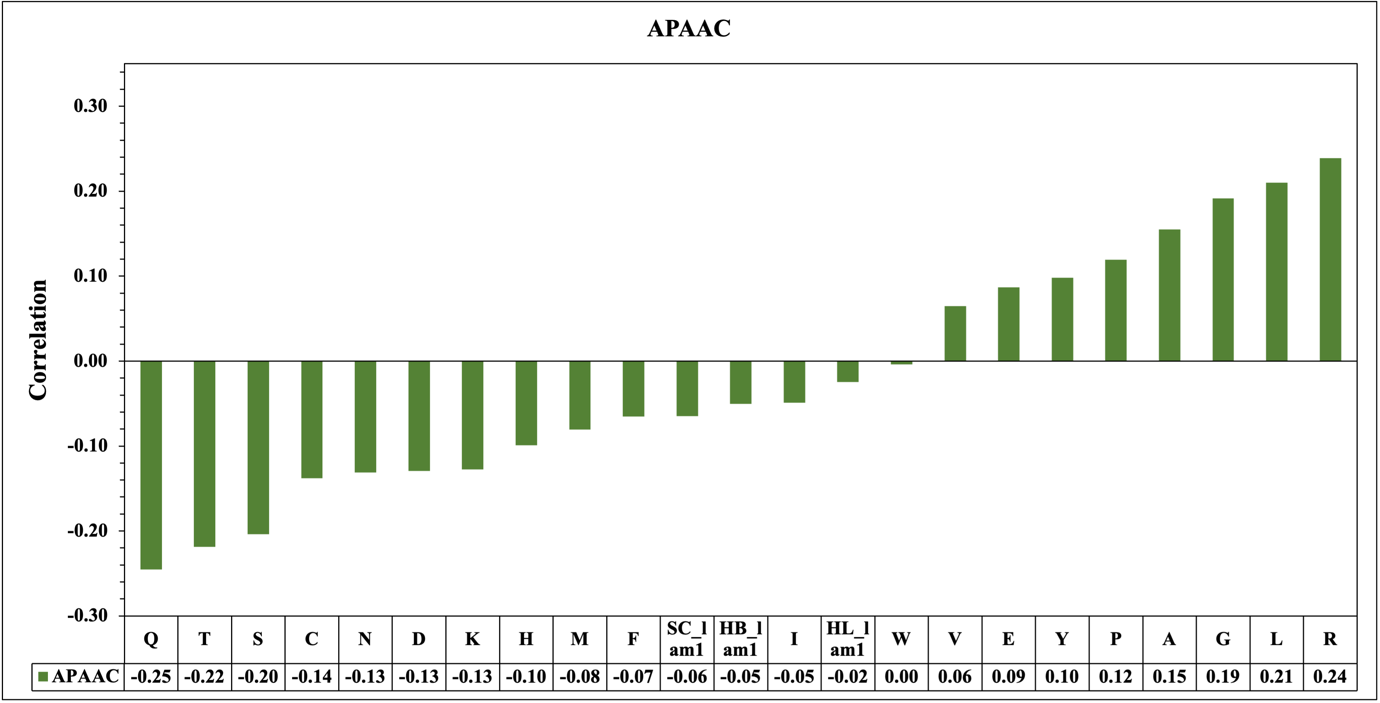

**Supplementary Figure S4: The correlation between the percent composition of APAAC residues and Tm**
